# Molecular basis for C-degron recognition by the SCF^Das1^ ubiquitin ligase

**DOI:** 10.64898/2026.05.13.724490

**Authors:** Maurice Bouchain, Julia K. Varga, Ka-Yiu Edwin Kong, Kay Hofmann, Ora Schueler-Furman, Anton Khmelinskii

## Abstract

Selective protein degradation by the ubiquitin-proteasome system frequently involves substrate recognition via short linear motifs known as degradation signals or degrons. Whereas degrons located at protein N-termini (N-degrons) have been extensively studied, our understanding of C-terminal degrons (C-degrons) is comparatively limited. Previously, we showed that the yeast SCF ubiquitin ligase and one of its substrate receptor subunits, the F-box protein Das1, target a broad range of C-degrons and implicated SCF^Das1^ in orphan quality control. Here, we sought to determine how Das1 recognizes its substrates. By combining *in vivo* competition assays with structural modeling and mutational analysis, we demonstrate that distinct C-degrons compete for a common site on Das1, indicating a shared mode of recognition. We identify a positively charged pocket within the Das1 leucine-rich repeat domain as the C-degron binding site. Three basic residues at the base of this pocket are essential for Das1 function, likely mediating electrostatic interactions with the C-terminal carboxyl group of the degron, while additional residues contribute to substrate specificity. Comparative analysis reveals that this pocket and its function are conserved across the Saccharomycetaceae family, supporting a conserved role for Das1 in protein quality control.

## Introduction

Proteostasis is maintained through a finely tuned balance between protein synthesis, folding, trafficking, assembly of complexes and degradation (Balch et al., 2008; Balchin et al., 2016). Central to the regulated removal of proteins is the ubiquitin-proteasome system (UPS), which selectively targets proteins for proteasomal degradation through their modification with ubiquitin. Substrate specificity within the UPS is largely dictated by E3 ubiquitin ligases, which recognize substrate features known as degradation signals or degrons (Zheng and Shabek, 2017). Whereas degrons can be either structural features or short linear motifs (SLiMs), the vast majority of degrons identified thus far are SLiMs (Ella et al., 2019; Zhang et al., 2025). Such degrons are frequently found in intrinsically disordered regions (IDRs) and are typically a few amino acids in length but can be as short as a single amino acid when located at protein N- and C-termini (N- and C-degrons, respectively) (Varshavsky, 2019; Timms and Koren, 2020; Sherpa et al., 2022).

A multitude of C-degron pathways was discovered in human cells, where different degron motifs are recognized mostly by interchangeable substrate receptors of CRL2 and CRL4 cullin-RING ubiquitin ligases (Koren et al., 2018; Lin et al., 2018; Zhang et al., 2025). Towards a comprehensive understanding of eukaryotic C-degron pathways, we recently performed an unbiased survey of C-degrons in budding yeast using multiplexed protein stability (MPS) profiling of peptide libraries (Kong et al., 2023b). This approach relies on the sfGFP-mCherry fluorescent timer (tFT) as a reporter of protein turnover. The mCherry/sfGFP ratio of fluorescence intensities in steady state anticorrelates with the turnover of tFT fusions, whereby a high mCherry/sfGFP ratio is indicative of a stable tFT fusion, and vice-versa (Khmelinskii et al., 2012; Fung et al., 2022). In MPS profiling, libraries of peptides fused to a tFT are expressed in yeast, followed by sorting of the yeast population into stability bins according to the tFT readout and deep DNA sequencing to identify the peptides present in each bin. Based on the distributions of sequencing reads, potential degrons are defined as sequences enriched in low stability bins (Kats et al., 2018; Kong et al., 2023a; Reinbold et al., 2023). Even though budding yeast lacks CRL2 ubiquitin ligases (as it lacks the Cul2 cullin) and the yeast Cul4/Rtt101 cullin of CRL4 complexes does not share significant homology with vertebrate cullins (Sarikas et al., 2011; Finley et al., 2012), we identified over 5000 potential C-degrons with MPS profiling of random peptide libraries and of the yeast C-terminome (Kong et al., 2023b). We determined that approximately 20% of these potential degrons are indeed C-degrons, and are targeted by the cullin-RING ubiquitin ligase CRL1/SCF. The SCF (Skp1, Cul1, F-box) is a conserved modular ubiquitin ligase that uses interchangeable F-box proteins with different specificities as substrate receptors (Zheng and Shabek, 2017). Even though there are 22 annotated F-box substrate receptors in yeast (Finley et al., 2012), virtually all the SCF-dependent degrons we identified appear to be short C-terminal motifs 4-5 amino acids in length that are recognized by a single F-box protein Das1 (Fig. S1a, b). Hasenjäger et al. independently arrived at similar conclusions (Hasenjäger et al., 2023).

Compared to CRL2 and CRL4 ubiquitin ligases targeting C-degrons in human cells, SCF^Das1^ appears to have a rather broad specificity. This notion is supported by two observations: the broad spectrum of random degrons and yeast C-terminal degrons targeted by Das1 and that most single amino acid substitutions at the 5 C-terminal positions have no impact on Das1 degrons (Kong et al., 2023b). The specificity of Das1 appears to be best reflected by a negative degron motif, dominated by disfavored amino acids such as charged amino acids, proline and tryptophan [DEKRPW] at the very C-terminus (position -1 hereafter), and negatively charged amino acids, glycine and proline [DEGP] at the preceding positions -5 to -2. Nevertheless, Das1 degrons are enriched in specific amino acids, including tryptophan (W) at position -5, lysine (K) at position -3, large hydrophobic amino acids isoleucine, leucine and methionine [ILM] at position -2 and cysteine and asparagine [CN] at position -1 (Fig. S1a). Moreover, while most Das1 degrons appear to be largely insensitive to mutations at position -5, we identified a set of degrons with an absolute requirement for a tryptophan at position -5 (referred to as W degrons hereafter) (Kong et al., 2023b). This suggests that Das1 targets at least two distinct classes of C-degrons, pointing towards one degenerate or multiple distinct degrons motifs. How Das1 recognizes such a broad range of degrons is unclear.

Here we set out to address this question and identify the substrate binding determinants in Das1. By combining structural modeling with mutagenesis and functional assays, we identified a single positively charged pocket in Das1 as the C-degron binding site. Our analysis indicates that this pocket and the broad specificity of Das1 towards C-degrons are conserved throughout the Saccharomycetaceae family of yeasts.

## Results

### Competition between Das1 degrons

Previously, we identified 931 Das1 degrons with MPS profiling of a library of 46152 random peptides 12 amino acid in length (tFT-X_12_ library) (Kong et al., 2023b). To understand how Das1 recognizes its substrates, we selected six Das1 degrons from this library: C(-1) degron (C at position -1), VN2 (N at position -1), Φ(-2) (a large hydrophobic residue at position -2), K(-3) (K at position -3), W(-5) (W at position -5) and IN2 (KIN motif at the very C-terminus) (Fig. 1a). We confirmed Das1-dependent turnover of the corresponding tFT-X_12_ constructs with a tFT colony assay, where strains expressing tFT-peptide fusions or tFT-tagged proteins are grown as ordered colony arrays, followed by measurements of colony fluorescence with a plate reader (Fig. 1b). Whereas most Das1 degrons are insensitive to mutations of position -5, mutating the tryptophan at position -5 in W degrons to any other amino acid completely abolishes Das1-dependent turnover of the corresponding tFT-X_12_ constructs (Kong et al., 2023b). We verified this behavior for the W(-5) degron (Fig. 1c). Whereas a tFT-W(-5) construct exhibited Das1-dependent turnover, a tFT-W(-5) variant with the tryptophan replaced by alanine (W(-5)A mutation) was completely stable. For comparison, mutating the last three residues to KIC makes a tFT-W(-5) variant insensitive to the W(-5)A mutation (Fig. 1c).

**Figure 1.**
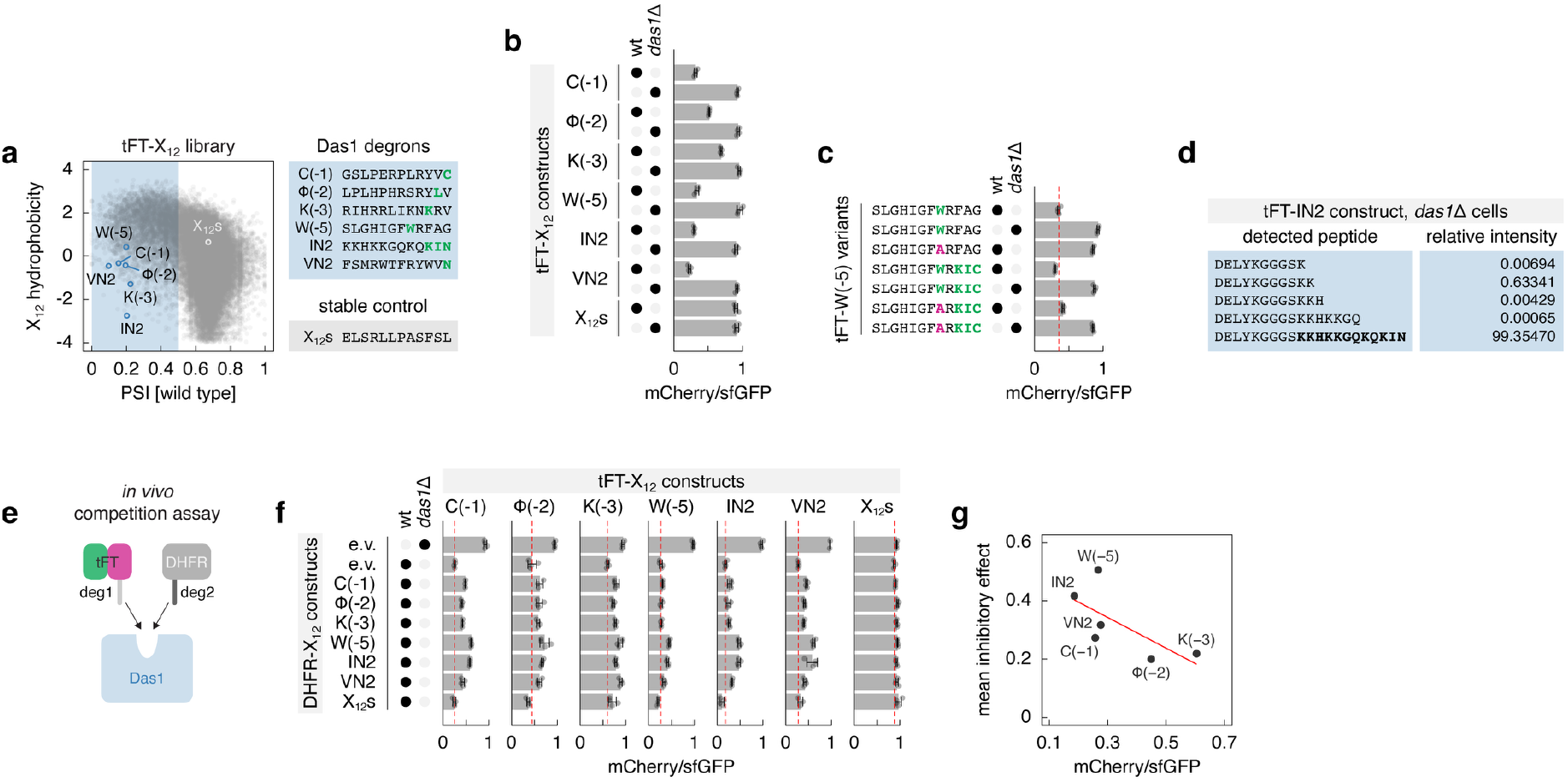
Competition between Das1 degrons. **a** – Correlation between peptide hydrophobicity (Kyte-Doolittle hydropathy scale) and protein stability index (PSI) in the tFT-X_12_ library analyzed with MPS profiling in (Kong et al., 2023b). Blue region marks putative degrons. Selected Das1 degrons and a control stable peptide (X_12_s) are indicated. **b, c** – mCherry/sfGFP ratios of colonies expressing tFT-X_12_ constructs (mean ± s.d., n = 4 biological replicates). wt, wild type. **d** – Mass spectrometry analysis of tFT-IN2 immunoprecipitated from *das1*Δ cells. Sequences and relative intensities of the identified C-terminal peptides. **e** – Cartoon of the *in vivo* competition assay with strains co-expressing different X_12_ peptides (deg1 and deg2) fused to a tFT or to DHFR. **f** – mCherry/sfGFP ratios of colonies co-expressing tFT-X_12_ and DHFR-X_12_ constructs (mean ± s.d., n = 4 biological replicates). e.v., empty vector; wt, wild type. **g** – Correlation between mean inhibitory effect of DHFR-X_12_ constructs in **f** and mCherry/sfGFP ratios of the corresponding tFT-X_12_ constructs in wild type cells transformed with an empty vector.

We asked whether degrons recognized by Das1 are post-translationally modified, e.g. by phosphorylation, proteolytic processing, intramolecular cyclization of glutamine or asparagine residues (which can generate C-terminal cyclic imide degrons (Ichikawa et al., 2022)) or α-amidating fragmentation (which can yield C-terminal amide degrons (Muhar et al., 2025)). Mass spectrometry analysis of the tFT-IN2 construct immunoprecipitated from cells lacking *DAS1* yielded no evidence of proteolytic processing or other C-terminal modifications, and almost exclusively identified a peptide containing the complete IN2 sequence (Fig. 1d). Similar analysis of tFT-C(-1) immunoprecipitates identified a peptide containing the complete C(-1) sequence and a series of shorter peptides lacking the last C-terminal residues (Fig. S1c). However, only the full length tFT-C(-1) construct exhibited Das1-dependent turnover in the tFT colony assay (Fig. S1d), arguing that shortened C(-1) sequences are not Das1 degrons. Together, this indicates that, at least in the case of the IN2 and C(-1) sequences, C-terminal modifications are unlikely to be involved in generating Das1 degrons.

Next, we sought to determine whether different C-degrons bind to the same or distinct sites on Das1. To distinguish between these possibilities, we established an *in vivo* competition assay based on co-expression of tFT-X_12_ and DHFR-X_12_ constructs. In case of competition for the same site on Das1, expression of non-fluorescent DHFR-X_12_ constructs should result in stabilization of tFT-X_12_ constructs, reflected by an increased mCherry/sfGFP ratio (Fig. 1e). In agreement with this expectation, expression of DHFR-C(-1) but not DHFR-X_12_s(where X_12_s is a peptide devoid of degrons that does not interact with Das1, Fig. 1a (Kong et al., 2023b)) stabilized tFT-C(-1) (Fig. 1f). Analysis of all possible construct combinations with the six Das1 degrons showed consistent competition between all degrons (Fig. 1f). Moreover, the mean inhibitory effect of DHFR-X_12_ constructs correlated with degron potency. For instance, more potent degrons corresponding to tFT-X_12_ constructs with lowest mCherry/sfGFP ratios were on average better competitive inhibitors in the assay (Fig. 1g). This suggests that, despite their substantial sequence diversity, different C-degrons likely bind to the same site on Das1.

### C-degron binding pocket in Das1

We sought to identify the degron-binding site on Das1. Das1 is a leucine-rich repeat (LRR) protein, with an F-box domain expected to interact with the Skp1 subunit of the SCF ubiquitin ligase. The uncharacterized F-box protein Ydr131c shares 37% of sequence similarity and 21% of sequence identity with Das1 and is predicted to adopt a similar fold (Fig. 2a). Previously, we found that Ydr131c does not target C-degrons recognized by Das1: tFT-X_12_ constructs with Das1-dependent turnover are not affected in cells lacking *YDR131C* and tFT-X_12_ constructs with partially reduced turnover in *das1*Δ cells are not further stabilized by deletion of *YDR131C* (Kong et al., 2023b). Despite their overall similarity, Ydr131c lacks three predicted intrinsically disordered regions (IDRs) present in Das1 (Fig. 2a). We thus asked whether these IDRs are required for targeting of C-degrons by Das1. We generated Das1 variants lacking the three IDRs, individually or combined, and a variant lacking the F-box (Das1-ΔF) as a control, with or without an N-terminal hemagglutinin (HA) epitope tag (Fig. 2b). N-terminally tagged HA-Das1 remained fully functional (Fig. S1e). With the exception of Das1-ΔF, all Das1 variants were expressed at levels comparable to the wild-type protein (Fig. 2c) and retained the ability to interact with Skp1 in a yeast two-hybrid (Y2H) assay (Fig. 2d). Using the tFT colony assay with six Das1 degrons, we found that deletion of the three IDRs had little to no impact on the ability of Das1 to target tFT-X_12_ constructs for degradation (Fig. 2e). This indicates that the predicted IDRs are not required for Das1 function and the degron binding determinants must be located elsewhere in Das1.

**Figure 2.**
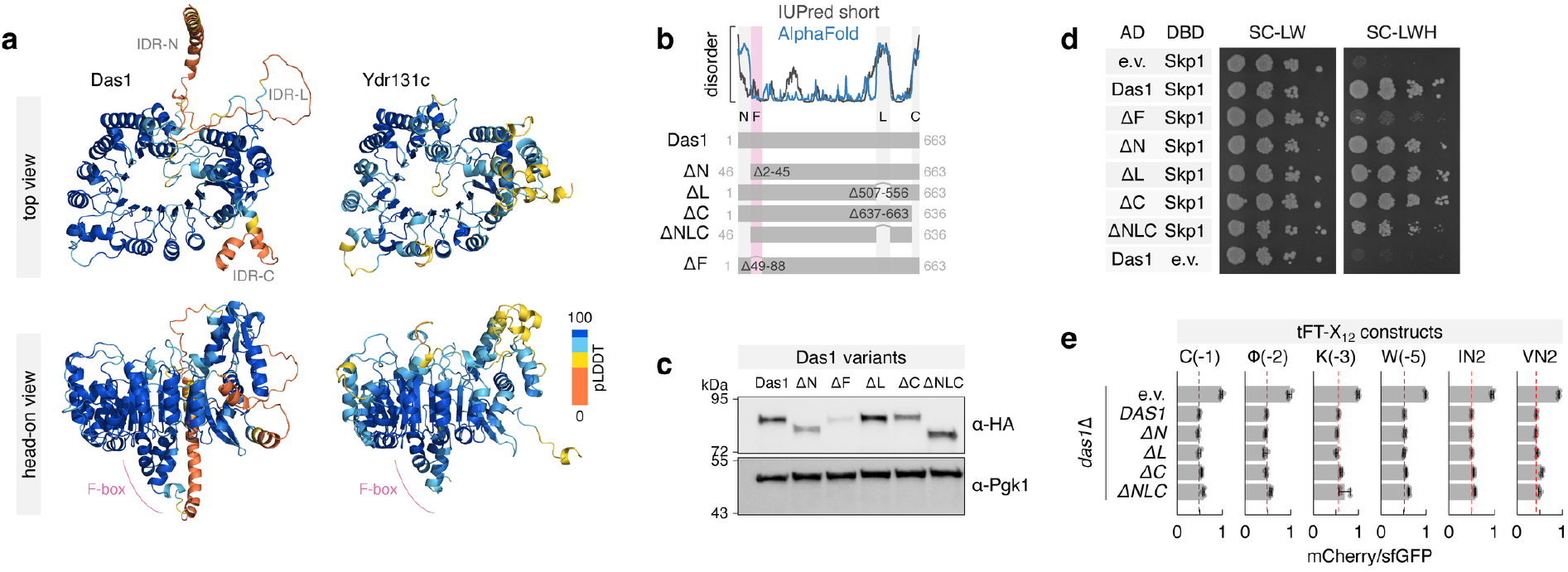
Functional analysis of the intrinsically disordered regions in Das1. **a** – AlphaFold models of Das1 and Ydr131c, color coded by predicted local distance difference test (pLDDT). Three intrinsically disordered regions in Das1 and the F-box motifs are marked. **b** – Cartoon of Das1 variants lacking the three predicted intrinsically disordered regions or the F-box motif. Top, disorder predictions. **c** – Levels of 3HA-tagged Das1 variants expressed from the *DAS1* promoter in a strain lacking endogenous Das1. Whole-cell extracts were separated by SDS-PAGE, followed by immunoblotting with antibodies against HA and Pgk1 as the loading control. **d** – Yeast two-hybrid interactions between Das1 variants fused to the activation domain (AD) and Skp1 fused to the DNA binding domain (DBD). Ten-fold serial dilutions on selective media lacking leucine and tryptophan (SC-LW, selection for the AD and DBD plasmids) or, in addition, lacking histidine (SC-LWH). Expression of His3, the two-hybrid reporter (Methods), only occurs upon interaction of AD and DBD fusion proteins. e.v., empty vector. **e** – mCherry/sfGFP ratios of colonies expressing tFT-X_12_ constructs (mean ± s.d., n = 4 biological replicates). Untagged Das1 variants were expressed from the *DAS1* promoter in strains lacking endogenous Das1. e.v., empty vector.

Next, we employed structural modeling with AlphaFold-multimer (Evans et al., 2021) to identify the potential degron-binding site in Das1. We selected a set of 50 Das1 degrons, 50 control (Das1-independent) degrons and 50 stable peptides from the tFT-X_12_ degron library that we had previously analyzed with MPS profiling (Kong et al., 2023b) (Fig. 3a). Das1 degrons were on average less hydrophobic than control degrons, and were enriched in cysteine and asparagine residues at position -1 and lysine at position -3 compared to both control degrons and stable peptides (Fig. S2a, b). We modeled each of these peptides with Das1 or Ydr131c for comparison and extracted the model confidence scores (Methods, Table S1). This analysis yielded on average higher confidence scores for models of Das1 with Das1 degrons compared to other degrons or stable peptides, and lower confidence for models of Ydr131c with all three groups of peptides (Fig. 3b), arguing for higher specificity of Das1 models with Das1 degrons. In these models, oriented with the F-box domain on the bottom face of the Das1 leucine-rich repeat, the peptide was typically found in a positively charged pocket on the top face, with its C-terminus at the bottom of the pocket (Fig. 3c).

**Figure 3.**
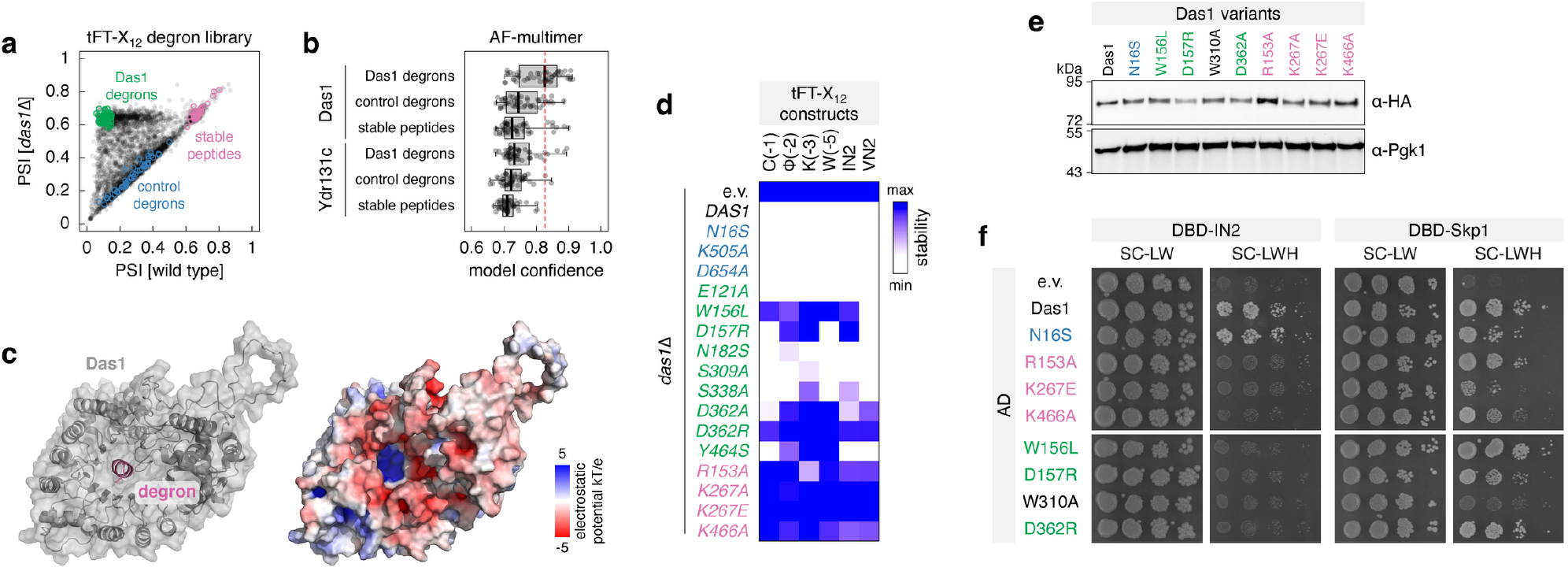
Substrate recognition determinants in Das1. **a** – Selection of X_12_ peptides for structural modeling. Protein stability index (PSI) in the tFT-X_12_ degron library analyzed with MPS profiling in wild type and *das1*Δ backgrounds in (Kong et al., 2023b). Three groups of selected peptides, with 50 peptides per group, are marked. **b** – AlphaFold-multimer models of X_12_ peptides from **a** with Das1 or Ydr131c. Distributions of model confidence scores. Red dashed line, median model confidence of Das1 models with 50 Das1 degrons. **c** – AlphaFold-multimer model of Das1 with the FTNLDYTKEKNN peptide (degron 49 in Fig. S2c). IDR-N and IDR-C are omitted for clarity. **d** – mCherry/sfGFP ratios of colonies expressing tFT-X_12_ constructs (mean ± s.d., n = 4 biological replicates). Untagged Das1 variants were expressed from the *DAS1* promoter in strains lacking endogenous Das1. e.v., empty vector. For each tFT-X_12_ construct, measurements were scaled as the change in mCherry/sfGFP readout relative to the strain expressing wild type Das1. e.v., empty vector. **e** – Levels of 3HA-tagged Das1 variants expressed from the *DAS1* promoter in a strain lacking endogenous Das1. **f** – Yeast two-hybrid interactions between Das1 variants fused to the activation domain (AD) and IN2 or Skp1 fused to the DNA binding domain (DBD). Interactions with DBD-IN2 performed in a *das1*Δ background.

To test the predicted C-degron binding pocket in Das1, we focused on the ten highest confidence models of Das1 with Das1 degrons. We identified 11 Das1 residues recurrently predicted to interact with Das1 degrons in these models: 3 basic residues (R153, K267 and K466) at the bottom of the pocket and 8 additional residues lining the pocket (Fig. S2c). Functional analysis of the corresponding single residue mutants in the tFT colony assay showed that the three basic residues R153, K267 and K466 are essential for Das1 function, whereas mutations in 5 of the other 8 predicted contact residues (W156, D157, S338, D362, Y464) affected the turnover of at least one tFT-X_12_ construct (Fig. 3d, Fig. S2d). For comparison, control mutations in the three IDRs (N16S, K505A and D654A) had no impact on the turnover of all tested constructs. Interestingly, none of the tested mutations in the 11 predicted contact residues or in 13 additional residues lining the pocket affected only the tFT-W(-5) construct (Fig. 3d, Fig. S2d, e), indicating that the binding mode of W degrons to Das1 remains to be defined. Importantly, pocket-disrupting mutations did not dramatically affect Das1 expression levels (Fig. 3e), with the exception of the D362R mutation (Fig. S2f). Moreover, all tested pocket mutants failed to interact with the IN2 peptide in the Y2H assay, while retaining the ability to interact with Skp1 (Fig. 3f). We conclude that C-degrons interact with a single pocket on Das1, whereby three positively charged residues at the bottom of the pocket likely engage in an electrostatic interaction with the negatively charged C-terminal carboxyl group of the degron, whereas additional residues lining the pocket are likely involved in degron-specific interactions.

### Das1 conservation and functions

Here we examined the conservation of Das1, its specificity and functions across evolution. Das1 orthologs could be readily identified in the Saccharomycotina subphylum (Groenewald et al., 2023), but only in three of the 16 families: Saccharomycetaceae, Saccharomycodaceae and Phaffomycetaceae (Table S2). We focused our analysis on the Saccharomycetaceae family, from the budding yeast *S. cerevisiae* to the filamentous fungus *Eremothecium gossypii*, which diverged approximately 100 million years ago (Dietrich et al., 2004). Whereas Das1 orthologs were identified in all species in the family, Ydr131c orthologs could be found in most but not all species (Fig. 4a). No Ydr131c orthologs could be identified in the four *Eremothecium* species (Table S2).

**Figure 4.**
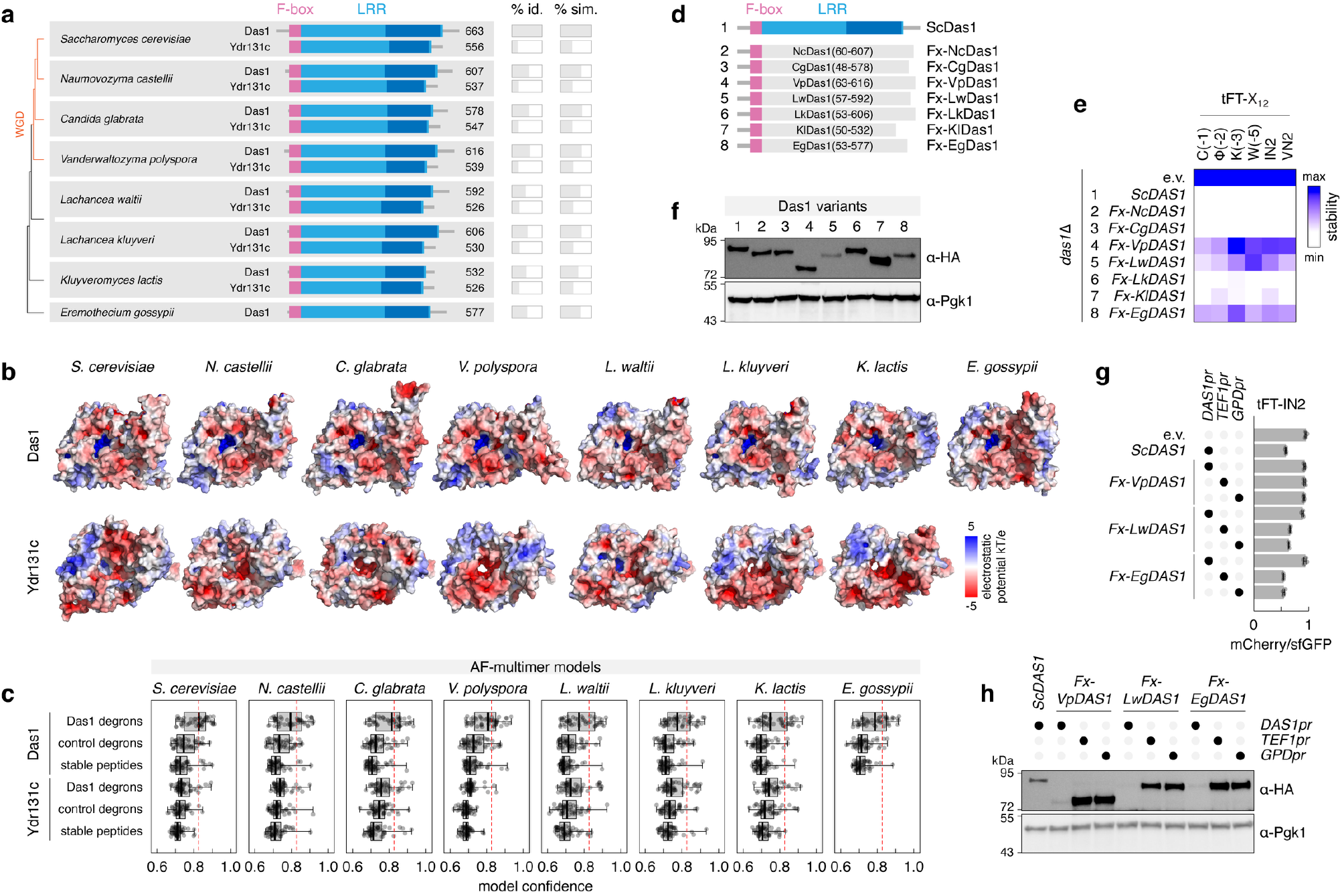
Das1 conservation across yeasts. **a** – Das1 and Ydr131c conservation across the Saccharomycetaceae family. The F-box motif and the leucine-rich repeat (LRR) are marked. Percentage of sequence identity and similarity (% id. and % sim., respectively) were calculated relative to *S. cerevisiae* Das1. **b** – AlphaFold models of Das1 homologs, color coded by surface charge. IDR-N and IDR-L in ScDas1, NcDas1, VpDas1, LwDas1, LkDas1 and IDR-N in EgDas1 are omitted for clarity. **c** – AlphaFold-multimer models of X_12_ peptides from Fig. 3a with Das1 homologs. Distributions of model confidence scores. Red dashed lines, median model confidence score of *S. cerevisiae* Das1 models with 50 Das1 degrons. **d** – Cartoon of Das1 chimeras combining the N-terminus (including the F-box) of *S. cerevisiae* Das1 with C-termini (from the start of the leucine-rich repeat) of Das1 homologs. **e** – mCherry/sfGFP ratios of colonies expressing tFT-X_12_ constructs (mean ± s.d., n = 4 biological replicates). Untagged Das1 chimeras were expressed from the *DAS1* promoter in strains lacking endogenous Das1. e.v., empty vector. For each tFT-X_12_ construct, measurements were scaled as the change in mCherry/sfGFP readout relative to the strain expressing wild type Das1. e.v., empty vector. **f** – Levels of 3HA-tagged Das1 chimeras expressed from the *DAS1* promoter in a strain lacking endogenous Das1. **g** – mCherry/sfGFP ratios of colonies expressing tFT-IN2 (mean ± s.d., n = 4 biological replicates). Untagged Das1 chimeras were expressed from the indicated promoters in a strain lacking endogenous Das1. e.v., empty vector. **h** – Levels of 3HA-tagged Das1 chimeras expressed from the indicated promoters in a strain lacking endogenous Das1.

Structural modeling and multiple sequence alignment showed the presence of a positively charged pocket, including the key basic residues R153, K267 and K466 implicated in binding C-degrons in the case of *S. cerevisiae* Das1, across all Das1 orthologs (Fig. 4b, Fig. S3). This suggests that both the structure and the function of this site might be evolutionarily conserved. In contrast, most Ydr131c orthologs lacked such a positively charged pocket (Fig. 4b). R153 was present only in *S. cerevisiae* Ydr131c, whereas K267 and K446 were not conserved across Ydr131c orthologs (Fig. S3). Structural modeling of different peptides (Fig. 3a) with Das1 or Ydr131c orthologs revealed a picture consistent with these observations: higher confidence for models of Das1 orthologs with Das1 degrons compared to other degrons or stable peptides, and lower confidence for models of Ydr131c orthologs with all three groups of peptides (Fig. 4c, Table S1). This further supports the notion that the broad specificity towards C-degrons is conserved among Das1 orthologs.

To experimentally assess this hypothesis, we analyzed the functionality of Das1 orthologs in the tFT colony assay by expressing them from the *S. cerevisiae DAS1* promoter in an *S. cerevisiae* strain lacking endogenous *DAS1*. To ensure proper interaction with *S. cerevisiae* Skp1, we used chimeric constructs in which the N-terminus of *S. cerevisiae* Das1, including the F-box motif, was fused to the leucine-rich repeat domain of each ortholog (Fig. 4d). Assays with tFT-X_12_ constructs carrying different Das1 degrons showed that most Das1 orthologs complemented the *DAS1* knockout and supported turnover of all tested constructs (Fig. 4e, Fig. S4a). However, the *V. polyspora, L. waltii* and *E. gossypii* Das1 chimeras appeared largely non-functional. As the *L. waltii* and *E. gossypii* Das1 chimeras were also poorly expressed (Fig. 4f), we asked whether reduced expression levels explain their apparent lack of function. Turnover of the tFT-IN2 construct was substantially increased upon expression of *L. waltii* and *E. gossypii* chimeras, but not the *V. polyspora* construct, from the strong constitutive *TEF1* and *GPD* promoters (Fig. 4g, h). The specificity of *V. polyspora* Das1 remains to be determined. Taken together, we conclude that the specificity of Das1 towards C-degrons is conserved in the Saccharomycetaceae family.

We also considered whether the *S. cerevisiae* Ydr131c could be engineered to recognize Das1 degrons. Based on the analysis of the C-degron binding pocket in Das1 (Fig. 3), we introduced mutations into Ydr131c to restore three residues critical for recognition of C-degrons: D157, K267 and K466 (Fig. S3). However, restoring these residues, alone or in combination, was not sufficient to confer Das1-like activity to Ydr131c expressed in cells lacking *DAS1* (Fig. S4b).

Evidence thus far suggests that Das1 is involved in protein quality control. One of the Das1 substrates is Hac1u, an aberrant protein resulting from translation of an unspliced mRNA encoding a key transcription factor in the unfolded protein response (Di Santo et al., 2016). In addition, Das1 targets for degradation the autophagy kinase Atg1 and the RNA polymerase I subunit Rpa12, specifically when these two proteins are produced in excess of their binding partners (i.e., when these two proteins are orphan) (Kong et al., 2023b). Mutations in the C-degron binding pocket of Das1, identified with artificial degrons (Fig. 3), impaired Das1-dependent turnover of overexpressed Atg1 and Rpa12 (Fig. 5a). This further strengthens the notion of this pocket as the substrate binding site in Das1.

**Figure 5.**
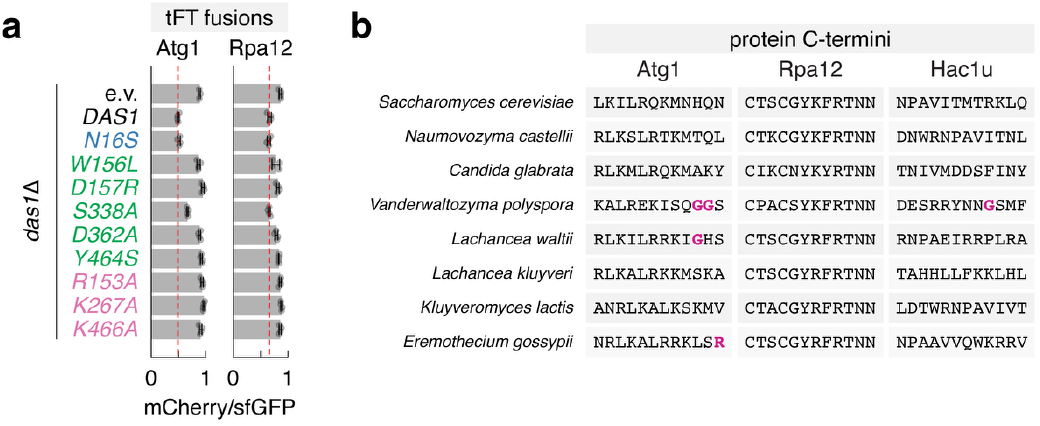
Conservation of Das1 substrates across yeasts. **a** – mCherry/sfGFP ratios of colonies expressing tFT-tagged full-length proteins from the strong constitutive *GPD* promoter (mean ± s.d., n = 4 independent clones per construct). **b** – C-termini of *S. cerevisiae* Das1 substrates across the Saccharomycetaceae family.

Considering the broad specificity of Das1, it seems reasonable to hypothesize that Das1 targets for degradation a wide range of thus far unidentified and potentially abnormal proteins. Here we examined +1 programmed translational frameshifting as a potential source of Das1 substrates. In *S. cerevisiae*, the Ty family of retrotransposons uses +1 programmed frameshifting in the expression of gag-pol fusion proteins (Voytas and Boeke, 1993; Farabaugh et al., 1993; Belcourt and Farabaugh, 1990; Clare et al., 1988). In addition, +1 programmed frameshifting is necessary for expression of Abp140, Est3, Oaz1 and Yfs1 proteins (Morris and Lundblad, 1997; Asakura et al., 1998; Palanimurugan et al., 2004; Ivanov et al., 2025). We asked whether C-termini of the corresponding short forms expected in the absence of a +1 programmed frameshift function as Das1 degrons. However, all tested constructs did not exhibit detectable Das1-depedent turnover and were stable in the tFT colony assay (Fig. S5b), indicating that Das1 is unlikely to be involved in protein quality control during +1 programmed frameshifting.

Finally, we asked whether the functions of Das1 might be evolutionarily conserved. Analysis of Atg1, Rpa12 and Hac1u sequences across the Saccharomycetaceae family showed that, in most species, the C-termini of these proteins are consistent with the negative degron motif targeted by *S. cerevisiae* Das1 (disfavored [DEKRPW] at position -1 of the C-degron and disfavored [DEGP] at positions -5 to -2 (Kong et al., 2023b)) (Fig. 5b). Therefore, we propose that Das1 involvement in protein quality control, potentially targeting Hac1u and orphan Atg1 and Rpa12 proteins for degradation, is prevalent across the Saccharomycetaceae family.

## Discussion

This work establishes the molecular basis for substrate recognition by SCF^Das1^, revealing that a single positively charged pocket within the leucine-rich repeat (LRR) domain of Das1 serves as the binding site for a diverse set of C-degrons. Three basic residues at the base of this pocket (R153, K267, and K466) are essential for function, most likely forming an electrostatic clamp around the free C-terminal carboxyl group of the degron, while flanking residues contribute degron-specific contacts.

These findings place SCF^Das1^ in comparative context with the C-degron pathways mediated by human CRL2 and CRL4 complexes. In humans, C-degron recognition appears to be distributed across a suite of interchangeable substrate receptors mainly of the CRL2 and CRL4 ubiquitin ligases – including KLHDC2, KLHDC3, KLHDC10, APPBP2, FEM1A/B/C, DCAF12 and TRPC4AP – each with narrow specificity for a particular C-terminal motif (GG, RG, PG and poly-alanine tails, RxxG, [RP], EE and Rxx, respectively) (Koren et al., 2018; Lin et al., 2018; Sherpa et al., 2022; Zhang et al., 2025). Structural studies of these receptors have revealed a recurring principle: a shallow generally positively-charged pocket in the receptor that engages the free C-terminal carboxyl group of a degron, while adjacent residues or pockets accommodate specific degron side chains to enforce sequence selectivity (Rusnac et al., 2018; Scott et al., 2024; Patil et al., 2023; Pla-Prats et al., 2023; Zhao et al., 2021; Yan et al., 2021; Chen et al., 2021). The positively charged pocket in Das1 is conceptually analogous, where the three essential basic residues likely constitute a carboxylate clamp. But Das1 achieves this through an LRR scaffold rather than the Kelch, tetratricopeptide, WD40 or ankyrin repeat folds used by CRL2 or CRL4 receptors (Zhang et al., 2025; Sherpa et al., 2022). This convergent solution to recognition of the C-terminal carboxyl group across phylogenetically and structurally distinct E3s underscores the chemical logic of C-degron surveillance. The failure to confer Das1-like activity to Ydr131c by restoring the three key basic residues alone implies that the broader electrostatic and geometric context of the LRR pocket, shaped by residues distributed across the repeat array, is collectively required for productive degron engagement, and that the binding site cannot be reduced to three anchor points. Notably, the C-terminal cyclic imide degrons recognized by CUL4^CRBN^ or the C-terminal amide degrons recognized by CUL1^FBXO31^ represent a further variation on this theme, where a post-translational modification, rather than a primary sequence motif with a C-terminal carboxyl group, generates a unique C-terminal feature recognized by the E3 (Ichikawa et al., 2022; Muhar et al., 2025), highlighting the chemical diversity of signals that E3s have evolved to detect at protein termini.

What distinguishes Das1 from its mammalian counterparts is its broad specificity. Whereas human C-degron receptors are highly selective for one or two C-terminal residues, Das1 tolerates extensive sequence variation across degron positions -1 through -5, with specificity defined more by a negative motif – especially exclusion of charged, proline, and tryptophan residues at position -1 – than by positive selection for a single chemical feature (Kong et al., 2023b). This permissiveness is not without precedent. For instance, the yeast nuclear quality control E3 San1 achieves broad substrate recognition through multiple degenerate binding sites that collectively detect exposed hydrophobic patches on misfolded proteins (Rosenbaum et al., 2011; Ibarra et al., 2021). However, Das1 operates through a distinct logic, recognizing a structural feature of the protein C-terminus itself (the free carboxyl group) rather than misfolding-induced exposure of internal sequences.

The broad specificity of Das1 is well suited to its described functions in protein quality control (Kong et al., 2023b; Hasenjäger et al., 2023; Di Santo et al., 2016). Our analysis suggests that this role of Das1 is likely conserved across yeasts, and across ∼100 million years of Saccharomycetaceae evolution in particular. However, considering its broad specificity and the few known substrates, it seems likely that further Das1 substrates, in particular under proteotoxic stress conditions, remains to be identified.

## Author Contributions

Conceptualization: A.K.; experimental investigation: all authors; writing-original draft: M.B. and A.K.; writing-review and editing: all authors; supervision: K.Y.E.K., O.S.F., A.K.; funding acquisition: K.H., O.S.F., A.K.

## Data Availability

AlphaFold-multimer model confidence scores for all models of the 150 peptides with Das1 or Ydr131c orthologs are provided as Table S1.

## Acknowledgements

We thank Joelle Strom and Katja Luck (IMB, Mainz, Germany) for help with scoring AlphaFold models. We thank Jia-Xuan Chen and the IMB Proteomics Core Facility for the mass spectrometry analysis; the IMB Bioinformatics Core Facility for the local infrastructure for structural modeling with AlphaFold; the IMB Media Lab and the IMB Protein Production facility for support with growth media and reagents. Funding of the Deutsche Forschungsgemeinschaft (DFG, German Research Foundation) supported the Orbitrap Astral system (P#524805621). J.K.V. was supported by a Marie Sklodowska-Curie European Training Network Grant #860517 (UBIMOTIF). Research in the O.S.F. lab was partially supported by the Israel Science Foundation (ISF) founded by the Israel Academy of Sciences and Humanities (grant no. 301/2021). This work was funded by the DFG (P#584323689 to A.K.).

## Methods

### Yeast strains and growth conditions

All yeast strains used in this work are listed in Table S3 and are derivatives of ESM356-1 or PJ69-4A. All plasmids used in this work are listed in Table S4. Yeast genome manipulations (gene deletions) were performed using PCR targeting and lithium acetate transformation (Gietz and Woods, 2002; Janke et al., 2004; Knop et al., 1999). Unless stated otherwise, yeast strains were grown at 30 °C in synthetic complete (SC) medium lacking histidine (SC-His) or uracil (SC-Ura) for plasmid selection, with 2% (w/v) glucose as carbon source.

### tFT colony assay

Fluorescence measurements of ordered colony arrays were performed as described (Kong et al., 2023b). Briefly, yeast strains were assembled in 1536-colony format on agar plates using a pinning robot (Rotor, Singer Instruments), with 4 biological replicates per tFT strain. For each biological replicate, 4 technical replicates of the sample strain, 8 technical replicates of a control strain without a tFT (non-fluorescent control) and 4 technical replicates of a reference strain expressing a stable tFT construct, were arranged next to each other in a 4×4 group. Such ordered colony arrays were typically grown for 24 h at 30°C before measuring colony fluorescence.

Fluorescence measurements were performed with a multimode microplate reader equipped with monochromators for precise selection of excitation and emission wavelengths (Spark, Tecan) and a custom temperature-controlled incubation chamber. Fluorescence intensities were measured as follows: mCherry with 586/10 nm excitation, 612/10 nm emission, optimal detector gain and 40 μs integration time; sfGFP with 488/10 nm excitation, 510/10 nm emission, optimal detector gain and 40 μs integration time. For each sample strain, fluorescence intensities were first corrected for background fluorescence by subtraction of the mean of neighboring non-fluorescent colonies, and then normalized by the mean of the neighboring reference colonies to correct for spatial effects. mCherry/sfGFP ratios were subsequently calculated for each technical replicate and summarized by the mean and standard deviation per biological replicate.

In the competition assay (Fig. 1e-g), the mean inhibitory effect of each DHFR-X_12_ construct was calculated as follows. First, for each tFT-X_12_ construct, all mCherry/sfGFP ratios were scaled between 0 and 1, by subtraction of the mean of the empty vector control in the wild type background, followed by normalization by the corrected mean of the empty vector control in the *das1*Δ background. Then, for each DHFR-X_12_ construct, the scaled mCherry/sfGFP ratios of the six tFT-X_12_ constructs (excluding the tFT-X_12_s control) were summarized by the mean.

### Yeast two-hybrid

All yeast two-hybrid experiments testing interactions between Das1 variants and Skp1 were performed with the PJ69-4A strain carrying a *HIS3* reporter (James et al., 1996). Interactions between Das1 variants and the IN2 peptide were tested in the yKEK476 strain, which was derived from PJ69-4A by knocking out *DAS1*. The strains were co-transformed with plasmids expressing Das1-AD variants and DBD-Skp1 or DBD-DHFR-IN2 fusions. Transformants were selected on SC plates lacking leucine and tryptophan (SC-LW). Selected transformants were grown in SC-LW overnight to saturation and serial ten-fold dilutions were spotted on SC-LW plates as a control for growth or on plates also lacking histidine (SC-LWH) to test for interactions between AD and DBD fusions. Growth on SC-LWH medium is only possible upon interaction between AD and DBD constructs, which activates *HIS3* expression. Plates were photographed (PhenoBooth imaging system, Singer Instruments) after 3 days of incubation at 30 °C.

### Immunoblotting

Approximately 5×10^7^ cells were harvested by centrifugation. Whole cell extracts were prepared by alkaline lysis followed by trichloroacetic acid precipitation (Knop et al., 1999). The precipitates were resuspended in high urea buffer (8 M urea, 200 mM Tris-HCl pH 6.8, 0.1 mM EDTA, 0.1% (w/v) bromophenol blue, 1.5% (w/v) DTT), followed by protein denaturation at 65°C for 15 min.

Whole cell extracts were then separated by SDS-PAGE, followed by immunoblotting with mouse primary antibodies against the HA epitope (dilution 1:2000, clone 12CA5, produced in-house) or Pgk1 (dilution 1:1000, 459250, Thermo Fisher Scientific) and a goat anti-mouse HRP-conjugated secondary antibody (dilution 1:5000, G-21040, Thermo Fisher Scientific). Membranes were developed after addition of a chemiluminescent horseradish peroxidase (HRP) substrate (SuperSignal West Pico PLUS, 34580, Thermo Fisher Scientific) and imaged using a ChemiDoc MP system (Bio-Rad).

### Mass spectrometry of tFT constructs

For immunoprecipitation, approximately 4×10^8^ cells were harvested by centrifugation, resuspended in IP buffer (25 mM Tris-HCl pH 8, 150 mM NaCl, 1% (v/v) Triton X-100, 1 mM EDTA, 0.5% [w/v] sodium deoxycholate, 0.1% [w/v] SDS and protease inhibitors (4693159001, Sigma-Aldrich)) and lysed by vortexing at 4°C with acid-washed glass beads. Lysates were clarified by centrifugation at 16000 to 21000 g. Clarified protein lysates were incubated with magnetic agarose beads coated with GFP nanobodies (Fridy et al., 2014). The bound proteins were eluted and denatured with elution buffer (25 mM Tris-HCl pH 8, 1% (w/v) SDS) at 90°C for 10 min with 1500 rpm shaking. Protein samples were then processed for mass spectrometry as detailed below.

Samples were processed using the SP3 approach (Hughes et al., 2019) [PMID: 30464214]. Briefly, proteins were reduced in 5 mM DTT, alkylated in 15 mM iodoacetamide in the dark and quenched in 5 mM DTT. Enzymatic protein digestion was performed using AspN or LysC protease at 37°C overnight. Following acidification by formic acid, the peptides were purified by solid phase extraction in a C_18_ StageTip format (Rappsilber et al., 2003). Peptides were separated via an in-house packed 45 cm analytical column (75 μm inner diameter; ReproSil-Pur 120 C_18_-AQ 1.9 μm silica particles, Dr. Maisch GmbH) on a Vanquish Neo UHPLC system (Thermo Fisher Scientific). Online reverse phase chromatography was performed through a 70 min non-linear gradient of 1.6-32% acetonitrile with 0.1% formic acid at a nanoflow rate of 300 nL/min. The eluted peptides were sprayed directly by electrospray ionization into an Orbitrap Astral mass spectrometer (Thermo Fisher Scientific). Mass spectrometry was conducted in data-dependent acquisition (DDA) mode using a top50 method with one full scan in the Orbitrap analyzer (scan range: 325 to 1300 m/z, resolution: 120000, target value: 3×10^6^, maximum injection time: 25 ms) followed by 50 fragment scans in the Astral analyzer via higher energy collision dissociation (HCD; normalized collision energy: 26%, scan range: 150 to 2000 m/z, target value: 1×10^4^, maximum injection time: 10 ms, isolation window: 1.4 m/z). Precursor ions of unassigned, +1 or higher than +6 charge state were rejected. Additionally, precursor ions already isolated for fragmentation were dynamically excluded for 15 s. Mass spectrometry raw files were processed using MaxQuant software (version 2.6.2.0 or 2.7.5.0) (Cox and Mann, 2008). MS/MS mass spectra were searched using Andromeda search engine (Cox et al., 2011) against a target-decoy database containing the forward and reverse protein sequences of the UniProt reference proteome for *S. cerevisiae* (release 2024_03 or 2024_04), the full-length reporter protein containing the C-terminal degron and its truncated versions with up to 11 amino acids deleted from its C-terminal end, and a default list of common contaminants. AspN or LysC specificity was assigned. Carbamidomethylation of cysteine was set as fixed modification. Methionine oxidation and protein N-terminal acetylation were chosen as variable modifications. The minimum peptide length was set to 7 amino acids. The “second peptide” search function was activated. The “match between runs” function was turned off. False discovery rate (FDR) was set to 1% at both peptide and protein levels. In the IN2 construct data, the detected peptide intensities were normalized by median-centering, based on the assumption that the overall peptide intensities were similar across the samples.

### Structural modeling with AlphaFold

Protein sequences were retrieved from the Saccharomyces Genome Database (SGD, https://yeastgenome.org) (Engel et al., 2025). Das1 and Ydr131c ortholog sequences were obtained from (Huerta-Cepas et al., 2014) and (Wapinski et al., 2007) or identified by sequence homology to *S. cerevisiae* Das1. The topology of the evolutionary tree was adapted from (Dujon, 2010). Multiple sequence alignment was performed using MUSCLE (Edgar, 2004).

Structural models of full length Das1 and Ydr131 orthologs were retrieved from the AlphaFold protein structure database (https://alphafold.ebi.ac.uk/) (Fleming et al., 2025). For clarity, N- and C-terminal IDRs were omitted for calculation of electrostatic potential surfaces as indicated (Fig. 3c, 4b). The model of the SCF^Das1^ complex, consisting of full length Cdc53, Skp1, Rbx1 and Das1ΔNLC (Fig. S1b), was generated with AlphaFold3 with default settings (Abramson et al., 2024).

Modeling of Das1 or Ydr131c orthologs from different species with 150 different 12 amino acid long peptides (Fig. 3b, 4c) was performed with AlphaFold multimer (Evans et al., 2021). Full length Das1 and Ydr131c ortholog sequences were used. For each model, five independent structural predictions were generated with a different seed per prediction to capture conformational variability and only the top prediction, selected based on the highest interface predicted template modeling (ipTM) and predicted template modeling (pTM) scores, was used for further analysis.

For downstream analysis, the top ten models of *S. cerevisiae* Das1 with Das1 degrons were identified by the model confidence score (0.8 x ipTM + 0.2 x pTM) (Fig. S2c). Das1 residues contacting the ten degrons were identified in Pymol (https://www.pymol.org) at a 4 Å distance threshold. Potential contact residues identified in at least three models were selected for mutational analysis.

## Figures

**Figure S1.**
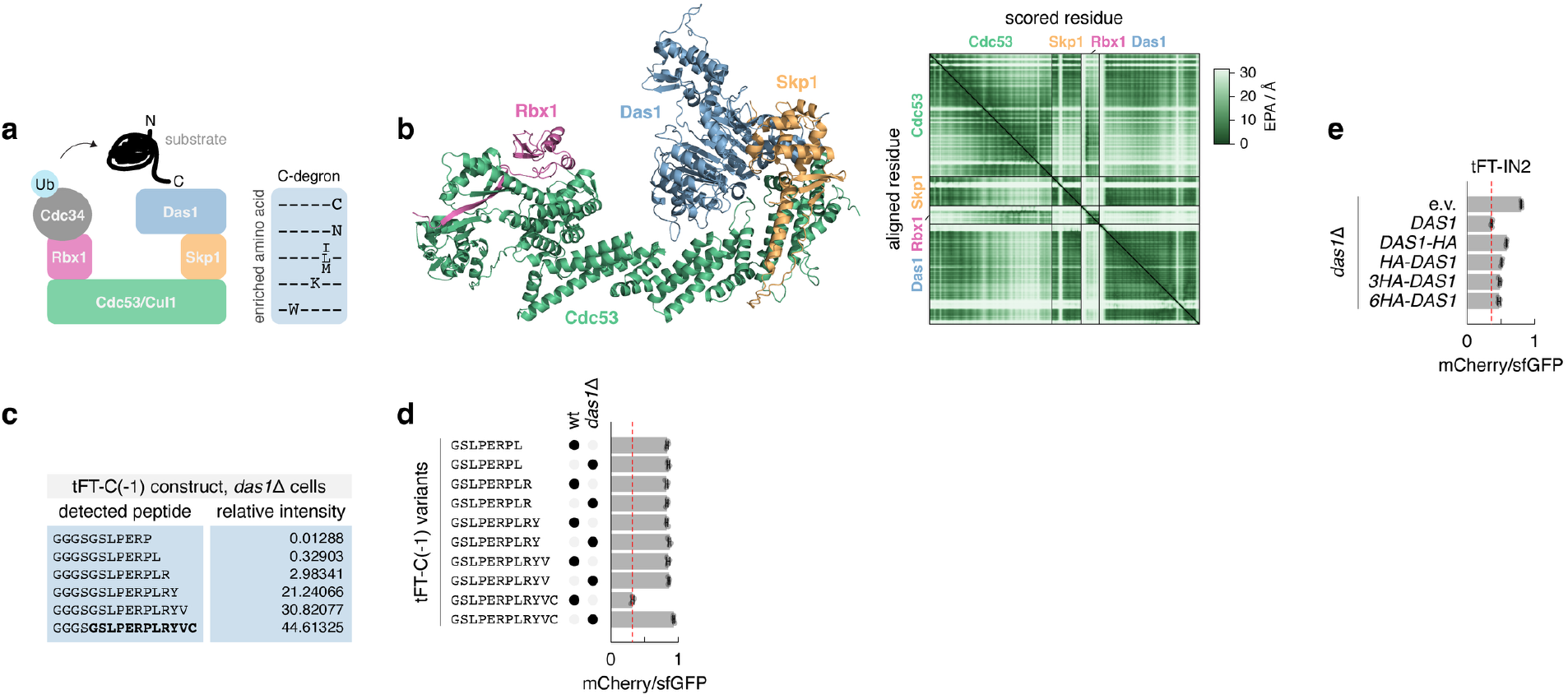
Functionality of HA-tagged Das1. **a** – Model of substrate recognition by SCF^Das1^ via C-degrons (left) and amino acids enriched in C-degrons targeted by Das1 (right). Adapted from (Kong et al., 2023b). **b** – AlphaFold-multimer model of the SCF-Das1 complex and the corresponding heatmap of expected predicted error (EPA). **c** – Mass spectrometry analysis of tFT-C(-1) immunoprecipitated from *das1*Δ cells. Sequences and relative intensities of the identified C-terminal peptides. **d** – mCherry/sfGFP ratios of colonies expressing tFT-X_12_ constructs (mean ± s.d., n = 4 biological replicates). wt, wild type. **e** – mCherry/sfGFP ratios of colonies expressing tFT-IN2 (mean ± s.d., n = 4 biological replicates). Das1 variants were expressed from the *DAS1* promoter in a strain lacking endogenous Das1. e.v., empty vector.

**Figure S2.**
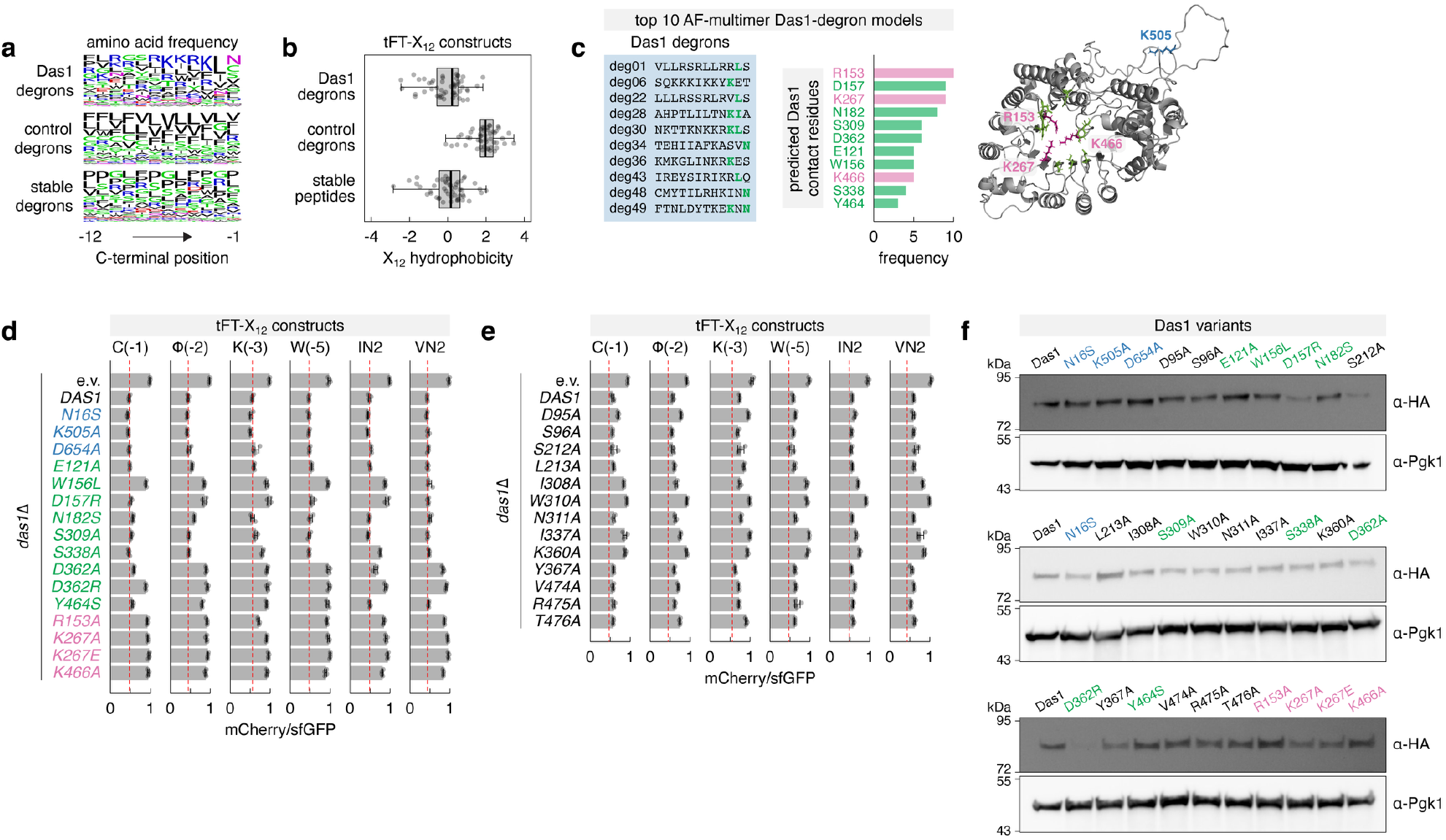
Substrate recognition determinants in Das1. **a, b** – Amino acid frequency and hydrophobicity (Kyte-Doolittle hydropathy scale) in the three groups of X_12_ peptides selected for structural modeling in Fig. 3a. **c** – Sequences of Das1 degrons (left) and Das1 residues recurrently predicted to contact C-degrons (center) in the top 10 AlphaFold-multimer models of Das1 with Das1 degrons in Fig. 3b. Right, AlphaFold model of Das1 with the predicted contact residues marked in magenta and green; IDR-N and IDR-C are omitted for clarity; K505 – a control residue not expected to interact with Das1 degrons. **d, e** – mCherry/sfGFP ratios of colonies expressing tFT-X_12_ constructs (mean ± s.d., n = 4 biological replicates). Untagged Das1 variants were expressed from the *DAS1* promoter in strains lacking endogenous Das1. e.v., empty vector. **f** – Levels of 3HA-tagged Das1 variants expressed from the *DAS1* promoter in a strain lacking endogenous Das1.

**Figure S3.**
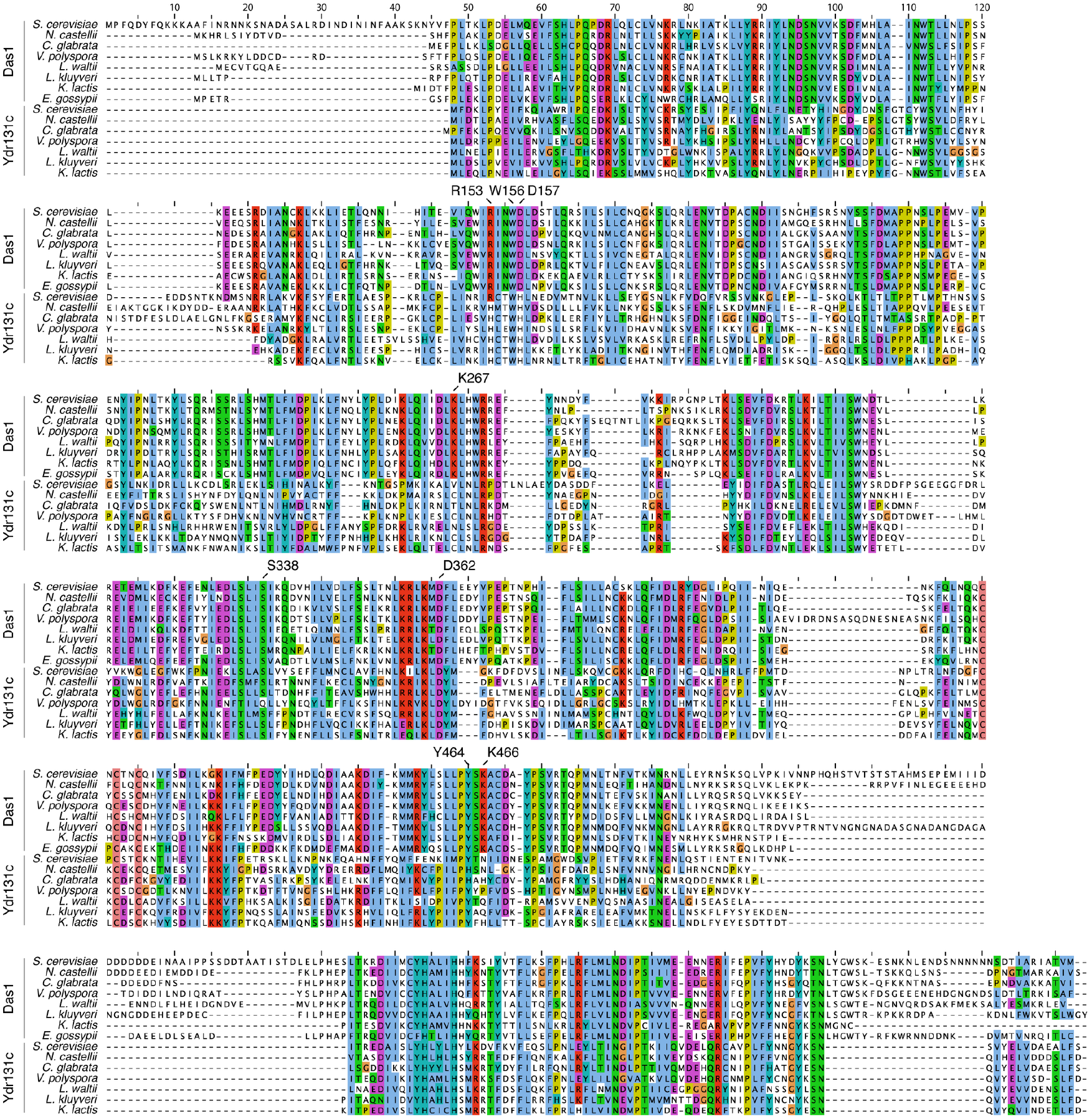
Das1 conservation across yeasts. Multiple sequence alignment of Das1 and Ydr131c homologs across the Saccharomycetaceae family. Key residues required for C-degron recognition by *S. cerevisiae* Das1 are marked above the alignment.

**Figure S4.**
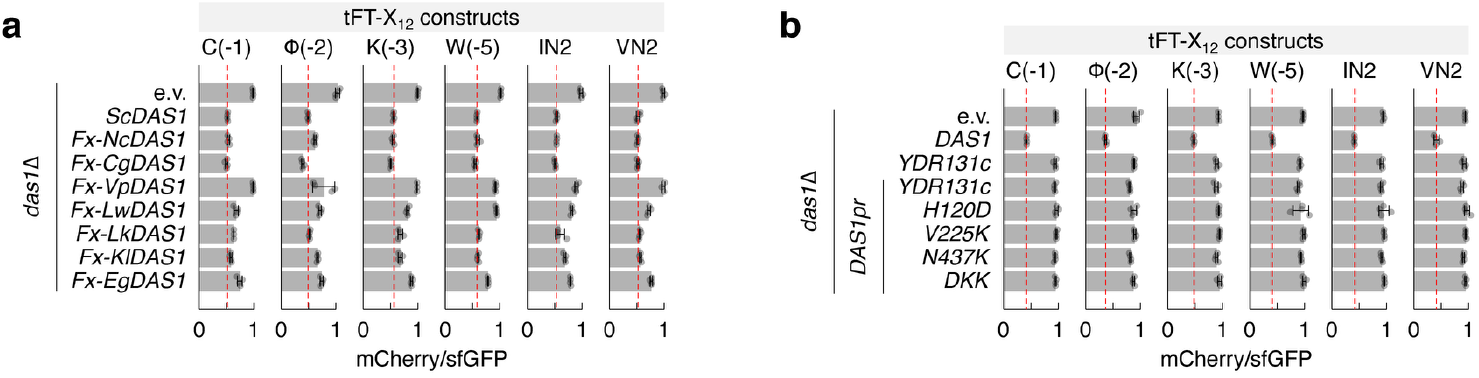
Specificity of Das1 homologs. **a** – mCherry/sfGFP ratios of colonies expressing tFT-X_12_ constructs (mean ± s.d., n = 4 biological replicates). Untagged Das1 chimeras were expressed from the *DAS1* promoter in strains lacking endogenous Das1. **b** – mCherry/sfGFP ratios of colonies expressing tFT-X_12_ constructs (mean ± s.d., n = 4 biological replicates). Untagged Ydr131c variants were expressed from the *YDR131C* or *DAS1* promoters in strains lacking endogenous Das1. e.v., empty vector.

**Figure S5.**
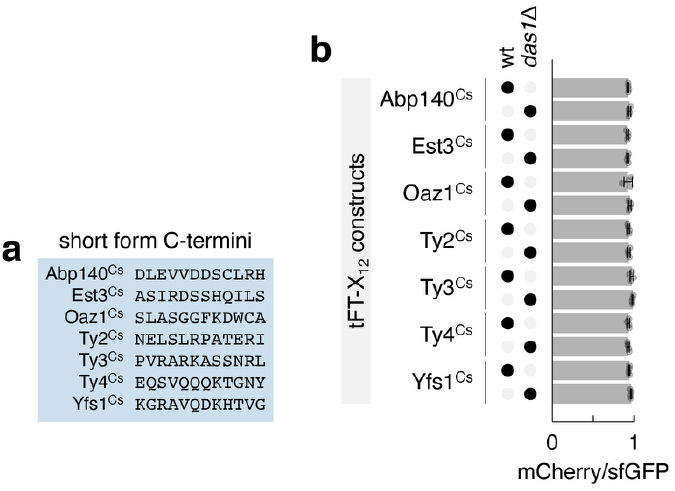
Das1 does not target C-termini resulting from failure in programmed frameshifting. **a** – C-termini of short protein forms expected from failure in programmed frameshifting during translation of the indicated mRNAs. **b** – mCherry/sfGFP ratios of colonies expressing tFT-X_12_ constructs (mean ± s.d., n = 4 biological replicates).

## Supplementary Tables

Supplementary Table 1 – Structural modeling of peptides with Das1 or Ydr131c.

Supplementary Table 2 – Das1 and Ydr131c orthologs across yeasts.

Supplementary Table 3 – Yeast strains used in this work.

Supplementary Table 4 – Plasmids used in this work.

